# Multiple infection of cells changes the dynamics of basic viral evolutionary processes

**DOI:** 10.1101/361766

**Authors:** Dominik Wodarz, David N. Levy, Natalia L. Komarova

## Abstract

The infection of cells by multiple copies of a given virus can impact virus evolution in a variety of ways, for example through recombination and reassortment, or through intra-cellular interactions among the viruses in a cell, such as complementation or interference. Surprisingly, multiple infection of cells can also influence some of the most basic evolutionary processes, which has not been studied so far. Here, we use computational models to explore how infection multiplicity affects the fixation probability of mutants, the rate of mutant generation, and the timing of mutant invasion. This is investigated for neutral, disadvantageous, and advantageous mutants. Among the results, we note surprising growth dynamics for neutral and disadvantageous mutants: Starting from a single mutant-infected cell, an initial growth phase is observed which is more characteristic of an advantageous mutant and is not observed in the absence of multiple infection. Therefore, in the short term, multiple infection increases the chances that neutral or disadvantageous mutants are present. Following this initial growth phase, however, the mutant dynamics enter a second phase that is driven by neutral drift or negative selection, respectively, which determines the long-term fixation probability of the mutant. Contrary to the short-term dynamics, the probability of mutant fixation, and thus existence, is lower in the presence compared to the absence of multiple infection, and declines with infection multiplicity. Hence, while infection multiplicity promotes mutant existence in the short term, it makes it less likely in the longer term. Understanding of these dynamics is essential for the investigation of more complex viral evolutionary processes, for which the dynamics described here for the basis. We demonstrate relevance to the interpretation of experiments in the context of published data on phage φ6 evolution at low and high multiplicities.

## Introduction

RNA viruses are characterized by very high mutation rates that are orders of magnitude faster than DNA viruses, due to the lack of proof-reading ability in RNA templated polymerases [1,2]. This, together with the typically large population sizes and rapid replication, promotes the generation of a large amount of genetic diversity that allows rapid adaptation to environmental challenges. The evolutionary dynamics of RNA viruses have been extensively studied in a variety of contexts [3-5]. Much of this work has viewed the virus genome as a solitary entity, where a specific gene in a given virus maps directly to its phenotype. It, however, has been demonstrated experimentally that genetically diverse viruses of the same species frequently co-habit a single cell, resulting in a variety of positive and negative interactions [6-12]. If different virus strains can interact in such ways, the “social structure” of the virus population becomes an important determinant of virus evolution, because interactions among viruses within cells can determine the response to selection and the level of genetic variation in the population [6,13-15]. Numerous examples of interactions among viruses in cells have been documented. Among positive interactions, viral complementation has been observed in several cases, leading to the persistence of inferior mutants [16-18]. Negative interactions are also possible, ranging from straightforward competitive interactions between viruses in a cell to the inhibition of the viral replicative potential [13,19,20]. Besides complementary and inhibitory interactions, different viruses coinfecting the same cell can exchange genetic information by recombination, for retroviruses such as HIV, and by reassortment for segmented viruses (e.g. influenza virus).

The effect of infection multiplicity (virus copies per cell) on evolutionary outcome has been examined in a variety of studies with different viruses[16,17,19-24]. Interesting results were obtained using the RNA phage φ6 [21,22]. For example, at high multiplicities of infection, defectors evolved that lowered the fitness of the phage population, which was not observed at low MOI. In a different study, viral diversity was found to be lower at high infection multiplicities, arguing that viral segmentation might have evolved for reasons other than the benefits of sex [23]. In the context of HIV-1, multiple infection has been shown to influence the latent state of integrated viruses [24]. That is, a latent virus in a cell can become activated through complementation when the cell is additionally infected by a productive virus. Overall, such work has shown that the effect of multiple infection on evolution is multi-factorial and complex.

While such complex social interactions certainly affect evolutionary dynamics in interesting ways that remain to be studied further, multiple infection of cells can also have the potential to influence basic viral evolutionary processes in simpler settings, which has so far remained under-explored. A solid understanding of the effect of multiplicity on the most basic evolutionary processes forms the underpinning for exploring more complicated scenarios. Here, we seek to contribute to this understanding with the help of evolutionary mathematical models. We investigate how infection multiplicity influences: (1) the fixation probability of a mutant virus starting from a single mutant-infected cell placed into a wild-type virus population at equilibrium, (2) the time until generation of the first mutant, and (3) the time to fixation in a model where mutant viruses are produced from wild-types with a defined rate. This is done in the context of neutral, disadvantageous, and advantageous mutants.

## The computational modeling framework

We study the evolutionary dynamics with a stochastic agent-based model because this allows for a natural formulation of the multiple infection process [25]. The model consists of N spots, which can be either empty, contain an uninfected cell, or contain an infected cell. Every time step, the system is randomly sampled N times, and the chosen spots are updated according to specific rules. If the chosen spot is empty, there is a probability L to produce an uninfected cell. If the sampled spot contains an uninfected cell, it can die with a probability D. If the sampled spot contains an infected cell, two events can happen. The cell can die with a probability A, and it can initiate an infection event with a probability B. If an infection event is initiated, a target spot is chosen randomly from the whole system. If that spot contains a susceptible cell, the infection event occurs, otherwise it is aborted. If the susceptible cell is an uninfected cell, it becomes infected with one virus. If the cell is already infected, its multiplicity is increased by one. The probability of an infected cell to die, as well as the probability to transmit a virus to another cell is assumed to be independent of infection multiplicity (different assumptions are explored below). The model assumes perfect mixing of viruses and cells.

The average dynamics of this system can be captured by ordinary differential equations. Denoting uninfected cells by y_0_ and cells infected by i viruses by y_i_, the equations are given as follows:

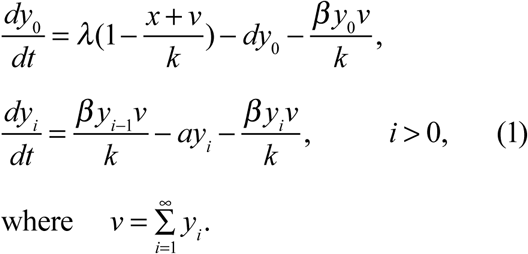

The variable v denotes the sum of all infected cells, which is proportional to the number of free viruses if free virus is in a quasi-steady state [26]. For numerical integration, this ODE formulation requires truncation at a maximum multiplicity, n, which needs to be large enough in computer simulations such that the population y_n_ remains negligible [25]. The virus establishes a persistent infection if its basic reproductive ratio, 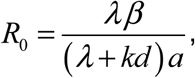 is greater than one. In this case, the dynamics converge to a stable equilibrium given by

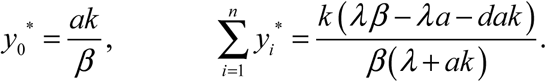

In the agent-based model, the populations will fluctuate around this equilibrium, due to the stochastic nature of the system, and the population of cells will be characterized by a given average infection multiplicity (Figure 1A & B).

**Figure 1.**
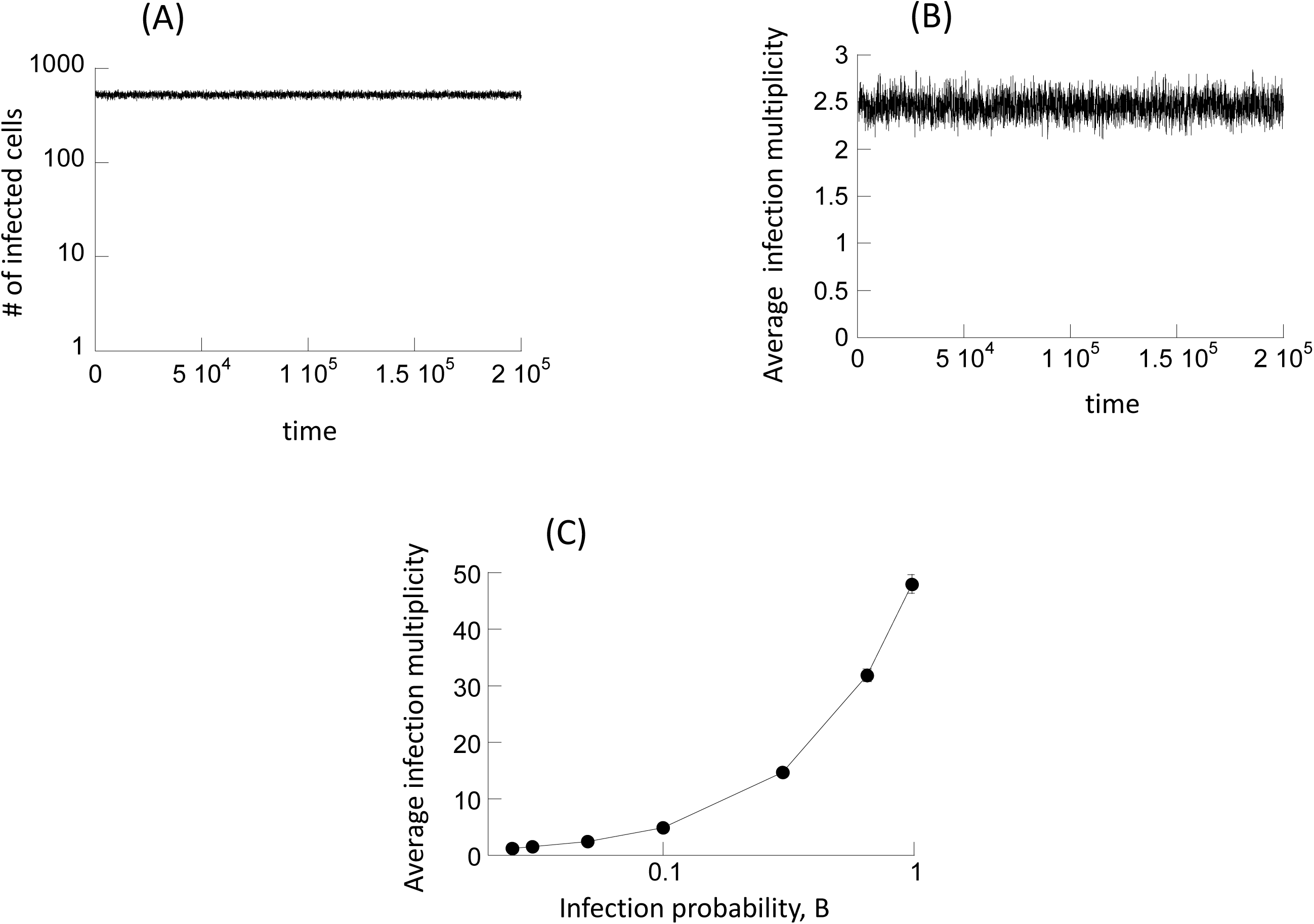
Basic properties of the computational modeling approach. The agent-based model is described in the text. (A) Over time, the number of infected cells converges towards and equilibrium value, around which the population fluctuates stochastically. A single typical run of the simulation is shown. (B) The average infection multiplicity across all infected cells also fluctuates around a steady state. Again, single typical simulation run is shown. (C) The average infection multiplicity is varied by changing the infection probability of the virus, B, as shown. The average multiplicity was determined by running the simulation repeatedly (10,000 runs), and taking the average value at a specific time point during the equilibrium phase of the dynamics. Standard deviations are plotted (almost not visible due to relatively small value). Base parameters are given as follows. B=0.025, A=0.02, L=1, D=0.01, N=900.

When a mutant virus is considered, there are two virus strains in the system that need to be tracked. The model follows cell populations that contain i copies of the wild-type virus, and j copies of the mutant virus. If a coinfected cell is chosen for infection, the virus strain to be transmitted is chosen randomly based on the fraction of the virus in the cell. Thus, the wild-type virus is chosen with a probability given by i/(i+j), and the mutant virus is chosen with probability j/(i+j) [25]. Again, the average dynamics of this system can be captured by ordinary differential equations. Denoting uninfected cells by y_00_ and cells infected with i copies of the wild-type virus and j copies of the mutant virus by y_ij_, the equations are given as follows:

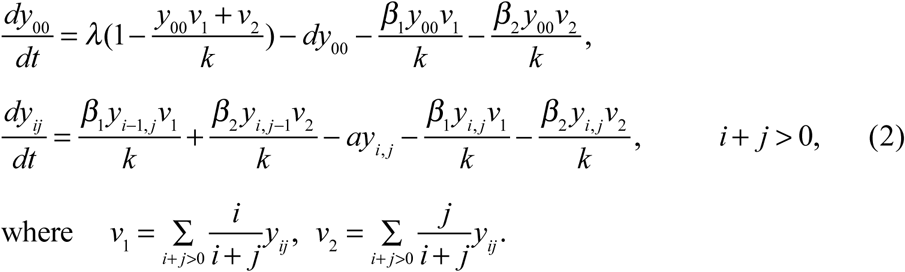

The variables v_1_ and v_2_ represent the sum of the fractions of the respective virus strains in the cell. This is proportional to the free virus populations if the rate of virus production is independent of multiplicity and if the virus is assumed to be in a quasi-steady state. The relative fitness of the two virus strains is determined by differences in the infection rates, β_1_ and β_2_. If these two rates are identical, the two virus strains are competitively neutral. For numerical integration, the system is truncated by only retaining the equations with i+j≤n, where n is sufficiently large.

## Varying the infection multiplicity

The goal of this work is to compare the evolutionary dynamics in settings where the multiplicity of infected cells is varied. This is achieved by increasing the infection probability B, because higher infection probabilities correlate with larger infection multiplicities at equilibrium, as shown in Figure 1C.

## Evolutionary dynamics of neutral mutants

We first consider the evolutionary spread of neutral mutants, i.e. the model parameters of the wild-type and mutant are identical. Different evolutionary endpoints will be considered in turn.

### Mutant fixation probability

We initialize the agent-based simulation by placing one cell with a single copy of the mutant virus (and no wild-type virus) into a population where the wild-type virus is present at equilibrium levels. The computer simulation was run repeatedly, and the fraction of simulations were determined that resulted in the fixation of the mutant. This is defined by the presence of the mutant virus, while the wild-type virus has gone extinct; realizations of the simulation in which both populations went extinct were not observed, and the simulation was set up to not count such events should they occur. The mutant fixation probability was determined for increasing infections rates, which correlate with higher infection multiplicities (Figure 1C). Systems with and without multiple infection were compared. In particular, to simulate the absence of multiple infection, infection events were aborted if the target cell was already infected with a virus. In the absence of multiple infection, the fixation probability of a neutral mutant is given by 1/N_cells_, where N_cells_ denotes the number of wt-infected cells at equilibrium before mutant introduction [27-29] (blue line, Figure 2A). This was verified by numerical simulations (not shown). The simulation results in the presence of multiple infection are shown in Figure 2A (black line, solid circles). For relatively low infection multiplicities (low infection probability, B), the observed fixation probability converges to the values in the absence of multiple infection, which is expected. The fixation probability, however, declines with increasing multiplicity, below the levels seen in the absence of multiple infection. Using the intuition from the theory of neutral evolution [27-29], in the presence of multiple infection, the fixation probability should be given by 1/N_viruses_, where N_viruses_ is the total number of viruses across all cells in the system; this is shown by the green line in Figure 2A. The observed fixation probability of the neutral mutant (black circles, Figure 2A), however, is significantly higher than this.

**Figure 2.**
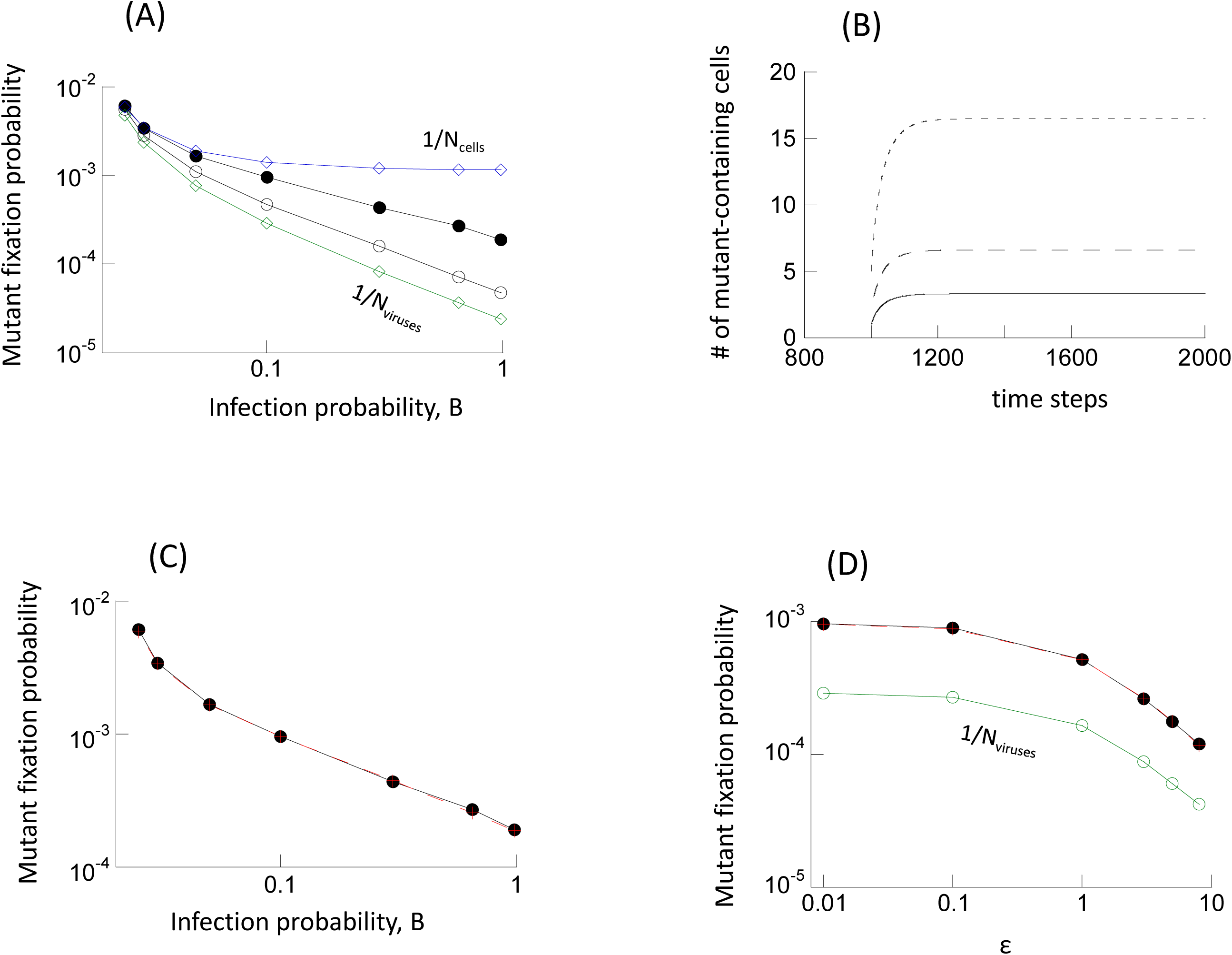
Evolutionary dynamics of neutral mutants. (A) Fixation probability as a function of the infection probability, B. Two theoretical bounds are shown by diamonds. The blue line with diamonds shows the fixation probability in the absence of multiple infection, given by 1/_Ncells_, where N_cells_ is the equilibrium number of infected cells before mutant introduction. The green line with diamonds shows 1/N_viruses_, where N_viruses_ is the equilibrium number of viruses across all cells before mutant introduction, and was hypothesized to be the theoretically expected fixation probability in the presence of multiple infection. The circles show results of two types of computer simulations. The black closed circles show the fixation probabilities in the computer simulation when one cell infected with one mutant virus is introduced into the system, where the wild-type virus population has equilibrated. The black open circles show the fixation probabilities in the computer simulation when one mutant virus is randomly placed into any of the available cells in the system where the wild-type virus population has equilibrated. Base parameters were: A=0.02, L=1, D=0.01, N=900. The number of simulation runs varied for different parameters due to different speeds of the computer simulation. For the black circles, the number of runs for increasing values of B were: 14154839, 15577853, 10415129, 18733054, 8590117, 5814742, 4518280. For open circles, the number of runs were: 29237576, 33491598, 24902642, 33231461,28297798, 22471381, 46938021. The trends described in the text are statistically significant, according to the Z test for two population proportions (very low p values, not shown). (B) Average dynamics of neutral mutants following introduction into a system at equilibrium, given by ODE model (2). The different lines depict simulations that start from different initial conditions. We observe first a phase of mutant spread, followed by convergence to a neutrally stable equilibrium. Parameters were: β=0.025, a=0.02, λ=1, d=0.01, k=900. (C) Successful theoretical prediction of the observed mutant fixation probability. The black circles show the observed fixation probabilities, which are the same as in panel A. The red crosses plot the values of Nneut/Nviruses, which accurately predict the observed fixation probabilities, as explained in the text. (D) Fixation probability of a neutral mutant in the agent based model where the rate of virus production is a saturating function of infection multiplicity. The fixation probability is shown as a function of the parameter ε, which determines how quickly saturation occurs. Higher values of ε correspond to a more pronounced increase in viral output with multiplicity. The green line again depicts the value of 1/N_viruses_. The red line plots the value of N_neut_/N_viruses_, which successfully predicts the observed fixation probabilities. Parameters were: B=0.025, A=0.02, L=1, D=0.01, N=900. The number of simulation runs for increasing values of e were: 119736073, 117908559, 104741112, 87608798, 75812069, 64365150. The trends described in the text are statistically significant, according to the Z test for two population proportions (very low p values, not shown).

The reason for this discrepancy is that there are two phases in the virus dynamics that contribute to this result. The average mutant dynamics are shown in Figure 2B, based on simulations of ordinary differential equation model (2). We observe that the population of mutant infected cells (which includes all cells that contain at least one mutant) initially grows, as if it were advantageous. This is followed by convergence towards a neutrally stable equilibrium (denoted by N_neut_, which depends on the initial mutant population size, Figure 2B). The initial growth phase, and hence, the initial advantage of the mutant, derives from the fact that in addition to uninfected cells, wt-infected cells also provide a target for new mutant infections. In contrast, new wt-infected cells can initially only be generated by viral entry into uninfected cells, since superinfection of wt-infected cells by more wt-virus does not result in the spread of the wild-type virus population. As the mutant spreads, this advantage diminishes and the dynamics enter the long-term neutral phase. This is because the mutant viruses become distributed among cells also containing wild-type virus and the initial asymmetry in growth dynamics vanishes. The initial advantageous phase of the dynamics accounts for the observed fixation probability that is higher than expected from the straightforward application of the neutral evolution argument. In fact, the number of mutant viruses (across all cells) at this neutral equilibrium, N_neut_, predicts the fixation probability, which is given by N_neut_/N_viruses_, where N_viruses_ is the total number of viruses before introduction of the mutant. This is shown in Figure 2C, where simulation results (black) are compared to the value of N_neut_/N_viruses_ (red). For this calculation, N_neut_ is determined by numerical integration of the ODEs.

In Figure 2A, the line with open circles depicts the results of additional simulations, which started from different initial conditions. Instead of introducing one cell that contains a single mutant virus, the mutant was placed into a randomly chosen (possibly infected) cell after the wild-type population had equilibrated. The fraction of runs in which mutants reached fixation was recorded. This corresponds to a scenario where the mutant was generated from the wild-type virus by mutational processes, and the fate of this mutant was followed for each realization of the simulation. Because mutant placement into a cell was probabilistic, in each simulation, the mutant virus was introduced into a different configuration, co-resident with different numbers of wild-type viruses within the cell. As seen in Figure 2A, the decline of the observed fixation probability of the neutral mutant with higher infection multiplicities is more pronounced in this case, and the fixation probability is closer to the value of 1/N_viruses_, but still higher. This makes intuitive sense, because the initial “advantageous” phase of the mutant dynamics is now less pronounced, due to intracellular competition of the first mutant virus with the wild-type.

### Time to appearance of first mutant

Another important evolutionary observable is the rate with which mutants are generated. This is explored here by quantifying the time it takes until the first mutant has been generated. To do this, we used a model that included mutational processes. When a wild-type virus was chosen for transmission to a new cell, it was assumed that a mutation occurred with a rate p_mut_. Biologically, this can correspond to mutations that occur upon production of the offspring virus, or that occur during the subsequent infection event (such as in retroviruses). For practical purposes, we chose a relatively high rate of p_mut_=3.5×10^-5^ per bp per generation, which is the mutation rate characteristic of HIV [30]. The dependence on infection multiplicity was explored in the same way as described above, by varying the infection probability. We found that for all infection rates, the time to first mutant generation is always faster in the presence compared to the absence of multiple infection (Figure 3A, compare black & blue line). Further, a higher infection multiplicity (infection rate) reduced the time at which the first mutant was generated (Figure 3A). This makes intuitive sense. A higher infection rate / multiplicity corresponds to more infection events, which in turn corresponds to more mutation events in this model.

**Figure 3.**
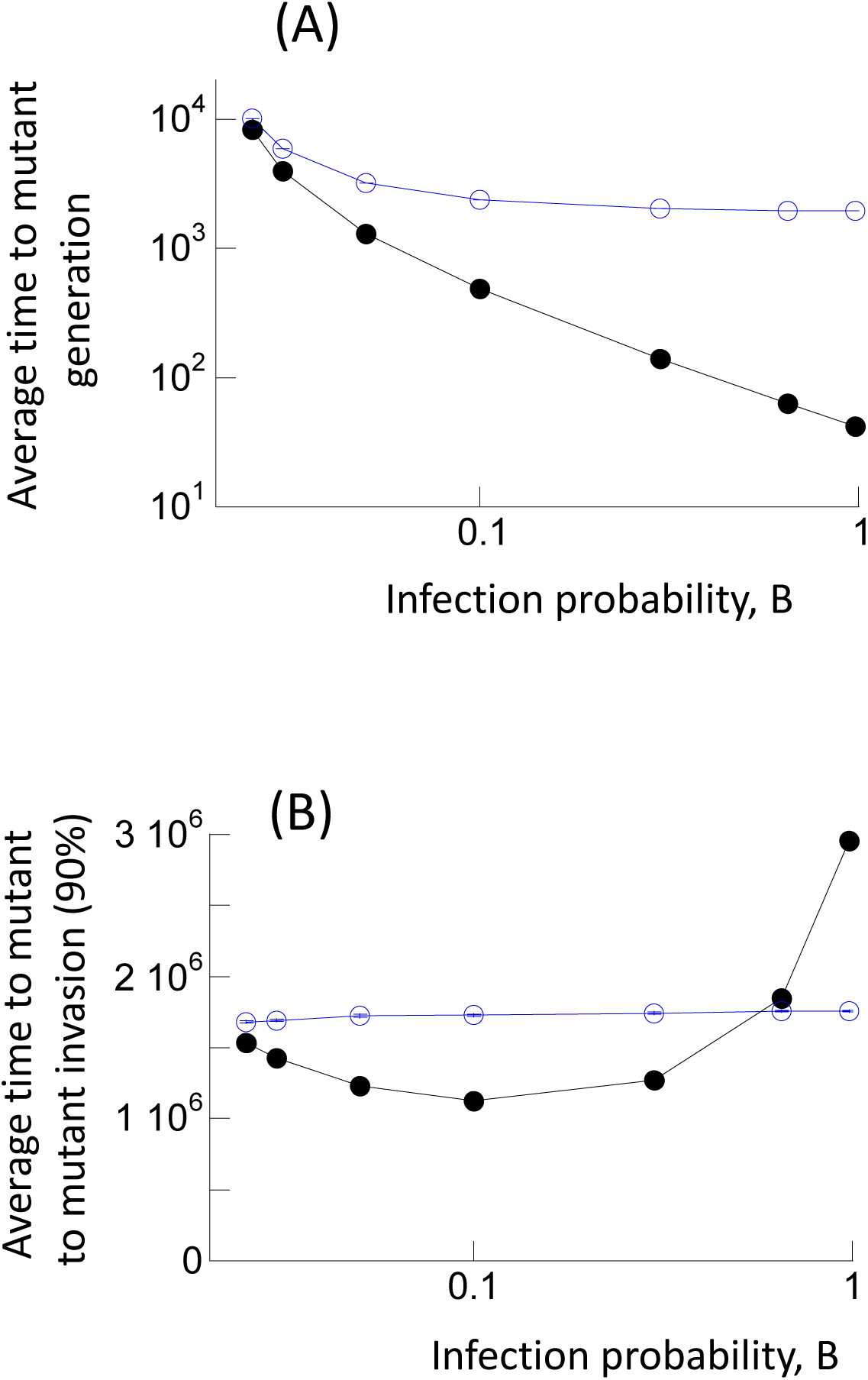
(A) Average time to generation of first mutant in the agent-based model with mutations. Black closed circles denote the simulation results in the presence of multiple infection, and blue open circles denote simulation results in the absence of multiple infection. Standard errors are shown, but are relatively small and hard to see. The number of simulation runs for increasing values of B for black circles are: 166137, 403110, 906346, 8000789, 8992529, 15656759, 19553451. For blue circles: 2159214, 4376870, 5980191, 10651321, 11229080, 10103400, 22652139. (B) Average time until the number of mutant-infected cells reached 90% of the whole infected cell population in the model with mutation and back-mutation (neutral mutants). The black closed circles show simulation results in the presence of multiple infection, the blue open circles show results without multiple infection. The simulation was started with the wild-type virus population at equilibrium. Parameters were chosen as follows: B=0.025, A=0.02, L=1, D=0.01, μ=3×10^-5^, N=900. Standard errors are shown, which, however, are very small and hard to see. For increasing values of B, we the number of simulations for the black circles was: 27629, 34858, 29688, 42050, 30574, 39744, 20570.. For blue circles: 34419, 39953, 29128, 38395, 34234, 72679, 64963. Trends of how multiple infection affects the plotted measures are statistically significant according to the 2-sample t-test (very low p values, not shown).

### Time to mutant fixation in a model with mutations

The above results indicate the existence of a tradeoff with respect to the effect of infection multiplicity. A higher infection multiplicity results in the more frequent generation of mutants. At the same time, however, it also leads to a lower probability of such mutants to invade and to fixate. The current section explores this tradeoff by using the model version with mutational processes and determining the time it takes for the mutant population to invade. In addition to wild-type giving rise to mutant viruses, however, we also need to account for back-mutations, since this counteracts the mutant expansion dynamics. In these simulations, the mutants are repeatedly generated (and eliminated at the same rate) and drift stochastically. Because of the occurrence of back-mutations, mutant fixation is not an absorbing state. To capture the effect of the tradeoff between increased mutant production and reduced invasion potential, we therefore recorded the time until the mutant reached 90% of the whole virus population for the first time (we refer to this event as “mutant invasion”). The results are shown by black circles in Figure 3B as a function of infection multiplicity. The corresponding results for simulations without multiple infection are shown in the blue line (Figure 3B). We find that multiplicity influences the time to mutant invasion in a non-monotonous way. For low viral infection rates (and hence low infection multiplicities), an increase in infection rate and multiplicity results in a reduced time to mutant invasion, which is below the time observed without multiple infection. As the infection rate and multiplicity are increased further, however, the time to mutant fixation becomes longer and rises above that observed in the absence of multiple infection (Figure 3B). Therefore, for moderate infection multiplicities, multiple infection speeds up mutant invasion. For higher infection multiplicities, multiple infection slows down mutant invasion.

## Disadvantageous mutants

Next we studied the evolutionary dynamics of slightly disadvantageous (0.05% fitness cost) mutants. The rules of the model are identical to those assumed for neutral mutants. In addition, once a virus was picked to infect a target cell, we assumed that this process failed with a probability 0.05% if this virus was a mutant, while it always succeeded if the selected virus was wild-type. In the absence of multiple infection, we numerically confirmed (not shown) that when one mutant-infected cell is introduced into a wild-type virus population at equilibrium, the fixation probability of the mutant is given by

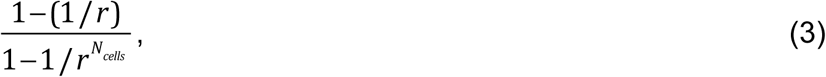

which is a formula derived from the Moran Process [31]. Here, r expresses the disadvantage of the mutant relative to the wild type, and N_cells_ denotes the number of wild-type infected cells at equilibrium before the mutant is introduced (see blue line, Figure 4A). In the context of multiple infection, the number of viruses rather than the number of cells should be the relevant population size, and hence by extension, the equivalent fixation probability would be given by

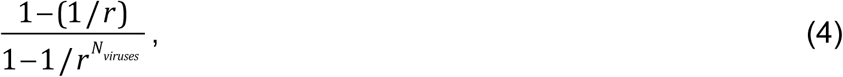

where N_viruses_ denotes the number of viruses across all infected cells (For reference, this is plotted by the green line in Figure 4A).

**Figure 4.**
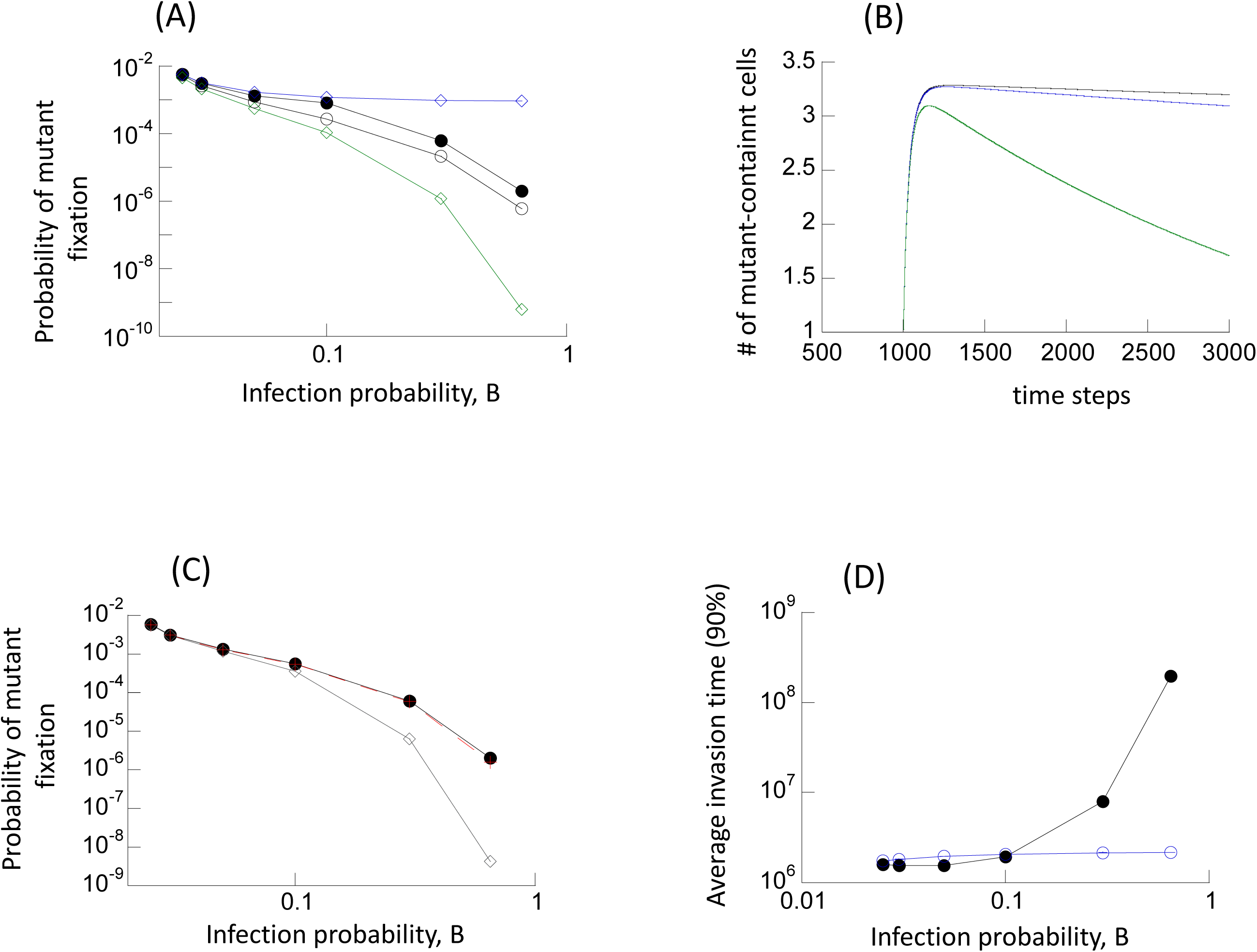
Evolutionary dynamics of disadvantageous mutants. (A) Fixation probability as a function of the infection probability, B. Two theoretical bounds are shown by diamonds. The blue line with diamonds shows the fixation probability in the absence of multiple infection as given by formula (3) derived from the Moran process. The green line shows the fixation probability according to formula (4) derived from the Moran process. The black closed circles show the fixation probabilities observed in the agent based simulation when one cell infected with one mutant virus is introduced into the system at equilibrium. The black open circles show the fixation probabilities observed in the agent based simulation when one mutant virus is randomly placed into any of the available cells in system at equilibrium. Base parameters were: B_1_=0.025, B_2_=rB_1_, A=0.02, L=1, D=0.01, μ=3×10^-5^, r=0.9995. The number of simulation runs for increasing values of B for the black closed circles were: 101317577, 112619340, 77957298, 37907585, 72473679, 47395056. For open black circles: 196598760, 225295595, 168947826, 227879849, 199753392, 277735577. Trends described in the text are statistically significant, according to the Z test for two population proportions (very low p values, not shown). (B) Average dynamics of disadvantageous mutants following introduction into a system at equilibrium, given by ODE model (2). We observe first a phase of mutant spread, followed by a decline towards extinction. Different lines depict different levels of mutant disadvantage. A larger disadvantage leads to a less pronounced initial spread phase, followed by a faster decline. Parameters were: μ_1_=0.025, β_2_=r β_1_ a=0.02, λ=1, d=0.01, k=900. The relative mutant fitness values were (from top to bottom) r=0.9995, r=0.999, and r=0.99. (C) Predicting the fixation probability of disadvantageous mutants. The black closed circles depict the same mutant fixation probabilities as in panel A, observed in agent based simulations that started with one cell containing one mutant virus with r=0.9995. The line with grey diamonds depicts the value of formula (5), assuming an initial mutant virus population size of N_neut_, and the relative mutant fitness disadvantage r=0.9995. This fails to accurately predict the observed fixation probability. The red line with crosses depicts the same formula (5), but using the composite mutant fitness value r’, defined in formula (6) in the text. This accurately predicts the observed fixation probability. (D) Average time until the number of infected cells containing the disadvantageous mutant reached 90% of the whole infected cell population, given by the agent-based model with mutations and back-mutations (black circles). The blue line depicts the same measure in the absence of multiple infection, determined by simulations of the agent-based model. Parameters were: B_1_=0.025, B_2_=rB_1_, A=0.02, L=1, D=0.01, μ=3×10^-5^, N=900, r=0.9995. Standard errors are shown but are relatively small and hard to see. The trends described in the text are statistically significant, according to the 2-sample t-test. The number of simulation results for increasing values of B for the black line was: 323339, 307142, 281610, 234979, 46647, 1338. For the blue line: 294547, 262278, 224520, 227618, 201695, 168677.

First, the simulations were started with one cell containing a single mutant virus being placed into a wild-type virus population at equilibrium (black closed circles, Figure 4A). Similar trends are observed compared to neutral mutants. The fixation probability of the disadvantageous mutant is found to be lower in the presence compared to the absence of multiple infection (Figure 4A, black closed circles and blue diamonds), and decreases with higher infection multiplicities. This decrease of the fixation probability with higher infection multiplicity is more pronounced than for neutral mutants. Nevertheless, the mutant fixation probability observed in the simulations is significantly higher than the one predicted by formula (4) (green line, Figure 4A). One reason for the higher fixation probability is the same as for neutral mutants. Despite its replicative disadvantage, the mutant initially enjoys an advantage over the wild-type virus, because in addition to un-infected cells it can also spread by entering wt-infected cells. Using the ODE model (2), this is shown in Figure 4B. The mutant cell population first rises. This is followed by a decline phase towards extinction, due to the assumed replicative disadvantage. The peak of the mutant dynamics curve is approximately the same as the neutral equilibrium that was observed for neutral mutants above (N_neut_). Hence, we hypothesized that the fixation probability of a disadvantageous mutant could be given by the Moran process formula assuming that the initial number of mutant viruses is given by N_neut_, i.e. by

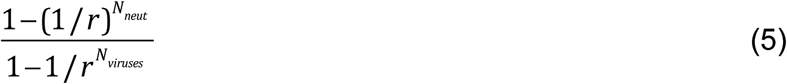

While this formula can predict the observed mutant fixation probability with reasonable accuracy for relatively low infection multiplicities (Figure 4C, grey diamonds), the observed fixation probability is significantly larger than this measure at higher multiplicities. The reason for this discrepancy seems to be that in the context of our model formulation, there are two levels at which mutant and wild-type viruses compete with each other: (i) Within a cell, a virus strain is picked for transmission with a probability given by the fraction of this strain in the cell. Hence the mutant is neutral with respect to the wild-type at this level. (ii) Between cells, the mutant is disadvantageous compared to the wild-type because it has a reduced probability to enter a new target cell (given by r<1). Therefore, the extent of the mutant fitness disadvantage is actually less than expressed by r, and the overall fitness of the mutant should be given by a value that lies between r and 1. The importance of this effect, however, should be influenced by the average multiplicity of the infected cells: If it is low, many cells contain either the mutant or the wild-type virus alone, and then the within-cell competition plays little role. In contrast, if the average infection multiplicity is high, most cells are likely to contain both mutant and wild-type virus, and the within-cell competition will play an important role. The overall fitness disadvantage of the mutant can thus be captured phenomenologically by an expression that places it between r and 1, weighed by the average infection multiplicity:

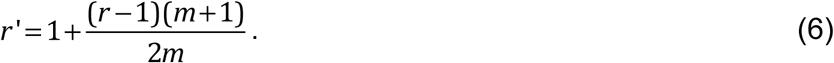

The parameter m denotes the average multiplicity among infected cells. If mutant fitness r’ is used in formula (5), we obtain a prediction that matches the fixation probability obtained in the computer simulation (red crosses superimposed on black circles in Figure 4C).

The curve with black open circles in Figure 4A again depicts the results of simulations in which the mutant was placed randomly in one of the available cells, and where the fate of the mutant was tracked. Because the first mutant virus now arises in a cell that could also contain wild-type viruses, the initial advantage of the mutant is less pronounced, as was the case for the neutral mutant. Hence, the mutant fixation probability is lower compared to that starting with a single mutant virus alone in a cell (closed black circles).

There is again a tradeoff between reduced fixation probabilities and the increased rates of mutant production with higher infection multiplicity (which is independent of mutant fitness). Again we recorded the time it takes for the mutant to reach 90% of the total virus population for the first time. The trend is similar to that for neutral mutants: at moderate infection multiplicities, multiple infection speeds up mutant invasion, but at higher multiplicities it slows down invasion (Figure 4D). The range of multiplicities over which mutant invasion is slower in the presence compared to the absence of multiple infection is wider for disadvantageous compared to neutral mutants (compare Figures 3B and 4D). Additionally, the extent to which multiple infection slows down mutant invasion is significantly stronger for disadvantageous mutants. Hence, multiple infection is more detrimental for mutant invasion for disadvantageous compared to neutral mutants.

## Advantageous mutants

Finally, we examined the evolutionary dynamics of advantageous mutants, assuming different degrees of mutant advantages (0.05%, 0.1%, 1%, Figure 5 A, B, C, resp.). The fitness advantage of the mutant was implemented similarly compared to the model for disadvantageous mutants: We assumed an overall infection probability that was 0.05%, 0.1%, and 1% higher than the value of the parameter B. When a mutant virus was selected to enter a target cell, this process was assumed to always succeed. When the wild-type virus was selected, there was a 0.05%, 0.1%, and 1% probability of failure. In this way, the wild-type virus had infection probability B, while the mutant virus had an overall higher infection probability. In the absence of multiple infection, the fixation probability is again given by formula (3) (see the blue lines in Figure 5(A-C)) derived from the Moran process, which we verified numerically (not shown). The parameter r>1 now measures the relative advantage of the mutant virus. As before, the green line shows formula (4), which is the Moran-process prediction for the fixation probability assuming that virus population size is given by the total number of viruses across all cells (rather than the number of infected cells).

**Figure 5.**
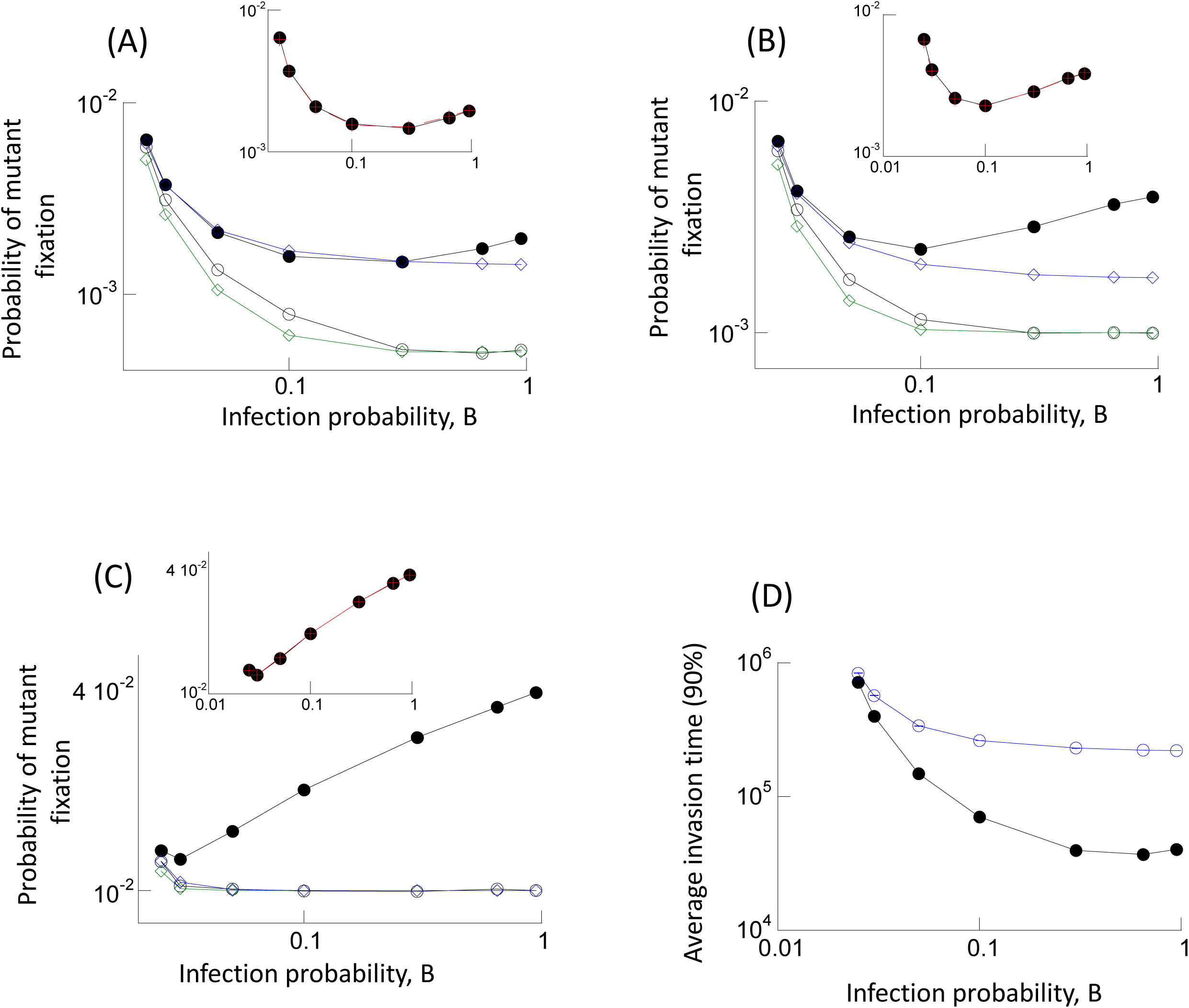
Evolutionary dynamics of advantageous mutants. (A) Fixation probability as a function of the infection probability, B. The blue line with diamonds shows the fixation probability in the absence of multiple infection, provided by formula (3). The green line shows the prediction of formula (4). The closed black circles show the fixation probabilities observed in the agent based simulation when one cell infected with one mutant virus is introduced into the system at equilibrium. The black open circles show the fixation probabilities observed in the computer simulation when one mutant virus is randomly placed into any of the available cells in system at equilibrium. The inset re-plots the observed fixation probability shown in closed black circles, and the red crosses depict the prediction given by formula (5) when the composite fitness value r’ is calculated according to formula (6), as described in the text. Parameters were: B_1_=0.025, B_2_=rB_1_, A=0.02, L=1, D=0.01, N=900, r=1.0005. The number of simulation results for black closed circles was: 4866352, 5371603, 3577510, 4091305, 2648860, 1691486, 1272341. For black open circles: 3935759, 4490230, 3313452, 4309990, 3470896, 4837538, 5286726. (B,C) Same simulations, but with larger mutant advantages, r=1.001 for B and r=1.01 for C. For B, the number of simulation runs for the closed black circles are; 7553368,6920249,6011459,5067733,3155326, 1939997, 1507107. For black open circles: 17417910, 19829787, 14557581, 18888450, 15088002, 11334612, 19407499. For C, closed black circles: 7519349, 6572315, 5445510, 4494334, 2831424, 1767559, 1362813. For C, black open circles; 854795, 773698, 702491, 681172, 547988, 407116, 333571. Trends described in the text are statistically significant, according to the Z test for two population proportions. (D) Average time until 90% of the infected cell population contain the advantageous mutant for the first time (black closed circles), based on the agent-based model with mutations and back-mutations, as a function of the infection probability. Standard errors are plotted, but are hard to see. The number of simulation runs are: 19839, 41663, 82828, 222263, 316597, 638422, 488754. The blue line depicts the result of equivalent simulations in the absence of multiple infection. Again, standard errors are too small to see, and the number of simulation runs are: 130825, 226054, 282790, 481975, 494045, 1080864, 998265. Parameters were: B_1_=0.025, B_2_=rB_1_, A=0.02, L=1, D=0.01, μ=3×10^-5^, N=900, r=1.01. The trends described in the text are statistically significant, according to the 2-sample t-test.

We again start from a single cell containing one mutant virus paced into a wild-type virus population at equilibrium, and determine the mutant fixation probabilities (shown with black closed circles in Figure 5(A-C) for different infection probabilities and hence multiplicities). The mutant fixation probability first declines with infection multiplicity (infection rate), and subsequently increases to levels that are larger than those observed without multiple infection. If the mutant has a larger advantage compared to the wild-type, this increase in the fixation probability is more pronounced (compare panels A, B & C of Figure 5). Hence, for sufficiently advantageous mutants, multiple infection largely increases the chances of mutant fixation.

The insets in panels (A-C) show that the fixation probability of the advantageous mutant is again accurately predicted by formula (5) derived from the Moran process, where the overall mutant fitness r’ is calculated by the empirical formula (6) (see red crosses superimposed on black circles). As before, this assumes that the initial number of mutant viruses is given by the neutral equilibrium (N_neut_, described in the context of neutral mutants above). This makes sense because the initial phase of mutant spread from the first cell (that only contains mutant virus) is similar for all mutant types as long as the fitness difference is not too large. Only after this initial virus dissemination does the competition between the two virus strains start to matter.

The open black circles show the results of simulations in which the mutant was generated randomly in any of the available cells once the wild-type virus population had equilibrated. Now, a drastically different trend is observed: the observed mutant fixation probability declines monotonically with infection multiplicity, as is also the case in the curve predicted by formula (4) (Figure 5(A-C), compare open circles and green line). The larger the extent of the mutant advantage, the closer the observed fixation probability is compared to the green line. In addition, we note that for more pronounced mutant advantages, the mutant fixation probability becomes largely independent of infection rate and hence multiplicity (Figure 5C). This indicates that for advantageous mutants, the nature of the initial conditions plays a very important role in determining how multiple infection influences the probability of mutant fixation.

As before, we also considered the physiologically more relevant scenario where a wild-type population at equilibrium is allowed to mutate with a probability p_mut_ per infection, thus repeatedly giving rise to the mutant virus. We find that the time to mutant invasion is always lower in the presence compared to the absence of multiple infection, and that an increase in multiplicity reduces the time until mutant invasion (Figure 5D). This follows from the above observations that (i) the advantageous mutant fixation probability shows a weak dependence on multiplicity if the mutant is placed randomly into any of the cells, and (ii) the rate of mutant generation is faster for higher infection multiplicity. Hence, in the context of advantageous mutants, multiple infection speeds up mutant invasion.

## Increased viral output in multiply infected cells

The analysis so far assumed that the amount of virus produced by infected cells during their life-spans is the same regardless of the infection multiplicity. This means that cellular factors limit the rate of virus production, and introduces an element of intracellular competition among the different virus strains. This section explores the effect of relaxing this assumption. The opposite extreme would be to assume that the rate of virus production is only driven by viral factors and that there is thus no intracellular competition among virus strains. In this case, the viral output from infected cells goes up with infection multiplicity. In particular, cells containing two viruses would produce twice as many offspring viruses, cells infected with three viruses would produce three times the amount of offspring virus, etc. This, however, would give rise to a positive feedback loop where higher multiplicity increases the rate of viral replication, which in turn increases the infection multiplicity. The biologically most reasonable assumption in this context would be that the rate of virus production is a saturating function of the number of viruses that are present in the cell. Hence, in the model, the probability for an infected cell to pass on the virus to a target cell is not given by B anymore, but by B(V)(1+ε)/(V+ε), where V denotes the total number of viruses in a cell. The larger the constant ε, the more the rate of virus production increases with multiplicity before converging to an asymptote. Thus, ε =0 corresponds to the case where viral output is independent of the multiplicity of infection, and ε → ∞ corresponds to the output increasing in an unlimited fashion with multiplicity. We investigated the fixation probability in the context of a neutral mutant. As the initial condition, we placed a single cell with one mutant virus into a wild-type population at equilibrium and recorded the mutant fixation probability as a function of the saturation constant ε. The results are shown in Figure 2D (black filled circles). For low values of ε, the fixation probability is close to the one observed for neutral mutants, where virus output was assumed independent of infection multiplicity. As the value of ε increases (more pronounced increase in viral output in multiply infected cells), the mutant fixation probability declines. This makes intuitive sense, because the average infection multiplicity in the cells rises with increasing values of ε. The green line (Figure 2D) again shows the reference value 1/N_viruses_. As before, the observed mutant fixation probability is significantly higher than the one predicted by neutral evolutionary theory (green line), for the same reason as given in the simpler versions of the model, where the rate of virus production was independent of multiplicity: the mutant dynamics first display a spread phase before the number of mutant viruses converges to a neutrally stable equilibrium (N_neut_). As in the simpler model in the previous sections, the fixation probability is again given by N_neut_/N_viruses_, as shown by the red crosses that are superimposed on the black circles in Figure 2D. Therefore, results described in the previous sections are not tied to the assumption that the rate of virus production is independent of multiplicity.

## Theory and data

An important aspect of theoretical work is relation to experimental data. While the dynamics of mutant fixation have not been studied in settings that vary the infection multiplicity, the number of mutants has been quantified in experiments where the bacteriophage φ6 was passaged under low and high infection multiplicity scenarios [23]. It was found that after 300 generations, genetic diversity was larger at low compared to high infection multiplicities, and that this difference was mostly due to the presence of mutations in non-coding regions of the genome, i.e. a result of neutral mutations. This suggested that processes occurring at high infection multiplicity (e.g. reassortment of genomic segments, sexual exchange), did not contribute to viral genetic diversity [23].

The models analyzed in our study made predictions about the average dynamics of neutral mutant viruses over time, which can be related to these experimental observations. In the presence of multiple infection, the dynamics were characterized by two phases: (i) An early growth phase was observed, where the dynamics resemble those of an advantageous mutant, which is not seen with neutral mutants in the absence of multiple infection. Hence, we expect that multiple infection promotes the spread of neutral mutants during this initial phase, which is counter to the experimental observations [23]. (ii) This initial phase is followed by convergence of the dynamics to a neutral equilibrium, the level of which predicts the long-term fixation / extinction probability of the mutant. Fixation is less likely with than without multiple infection, and declines with higher infection multiplicities. Stated in another way, the mutant virus is more likely to go extinct in the presence compared to the absence of multiple infection, and higher multiplicities further promote mutant extinction. Therefore, during this longer term phase, the number of neutral viruses is predicted to be larger at low compared to high multiplicities. This is in agreement with the experimental data on the evolution of phage φ6 [23].

Another complication in the interpretation of the experimental data concerns the experimental measure under consideration. The model suggests that different results can be obtained about the average number of mutants at low and high infection multiplicities depending on whether the number of mutant-infected cells are counted, or whether the amount of free virus is compared. This is demonstrated with computer simulations in Figure 6. As initial conditions, a model simulation with wild-type virus only was allowed to equilibrate, and 10% of the infected cell population was sampled to start a new growth phase. A small number of mutant viruses was added to this pool of cells and the resulting growth curves were recorded. This might mimic a virus passage, which was part of the experiments performed by Dennehy et al [23]. Many repeats of such runs were performed, and the average population sizes, as well as standard errors are plotted over time in Figure 6. Figure 6A shows that if the number of mutant-infected cells is compared, the number is larger in the presence compared to the absence of multiple infection. In contrast, Figure 6B shows the opposite if a measure proportional to the free virus population is compared. Even though more mutant-infected cells are predicted in the presence of multiple infection, if the mutant virus is significantly diluted by wild-type copies within those cells, fewer mutant free viruses will be observed with multiple infection. The reason is the assumption that the rate of mutant virus production is proportional to the fraction of the mutant in the cell.

**Figure 6.**
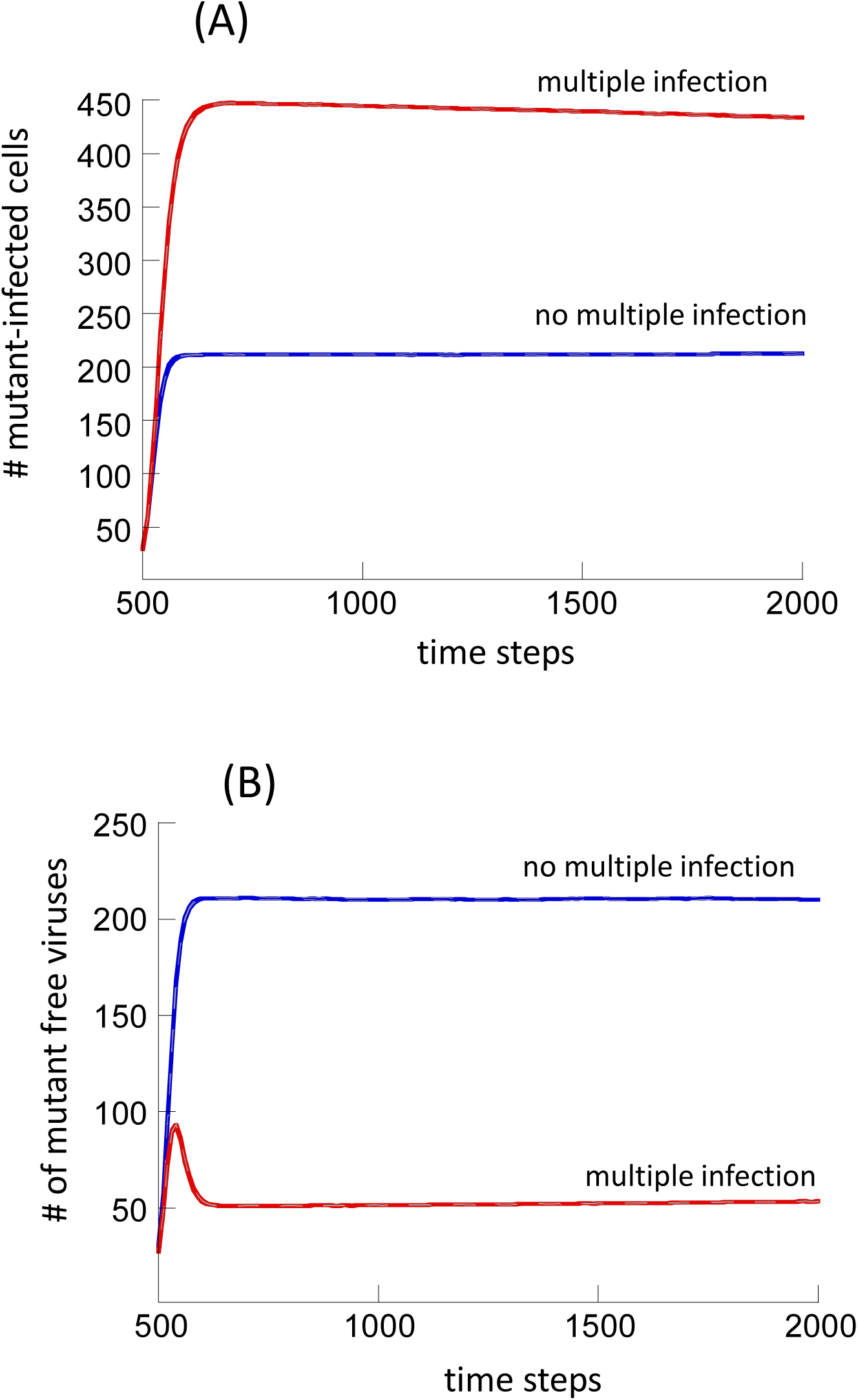
Average mutant dynamics in the presence (red) and absence (blue) of multiple infection, based on repeated realizations (100,000) of the agent-based model without mutational processes. The grey dashed lines depict the standard errors. The simulations were started with wild-type virus only, until the system equilibrated. Then, 10% of the wild-type-infected cells were randomly selected, and renewed growth was simulated, together with a minority population of mutants (30% of the wild-type population). This mimics the basic virus passage procedures in phage experiments reported by Dennehy et al [23]. (A) The number of mutant-infected cells is plotted. (B) The sum of the mutant fractions across all infected cells is plotted, which is proportional to the amount of free virus. Parameters were: B=0.025, A=0.02, L=1, D=0.01, N=900.

In summary, the models have identified two factors that can impact whether mutant spread is intensified in the presence or absence of multiple infection. The timing after mutant introduction can determine the result, and so can the particular measure of mutant spread. These complexities are important to keep in mind when interpreting experimental data.

## Discussion and Conclusion

We used computational models to investigate the spread dynamics of mutant viruses in the presence of multiple infection, assuming relatively simple settings where no viral complementation, inhibition, or recombination / reassortment occurred. Nevertheless, the dynamics were found to be complex. An interesting aspect concerns neutral and disadvantageous virus mutants. During the initial stages of the dynamics, the mutant population enjoys growth instead of drifting, similar to an advantageous mutant. The reason is than an initial asymmetry confers an advantage to the mutant relative to the wild-type virus. Specifically, in a virus population that contains almost only wild-types, new wt-infected cells can only be generated by viral entry into uninfected cells. On the other hand, new mutant-infected cells can be generated both by entry into uninfected cells, and by entry into wild-type infected cells. This can be tested experimentally by labeling viruses with two different fluorescent colors and introducing a minority population of one color (the “mutant”) into a population that contains a relatively large number of uninfected cells, as well as cells infected with the virus labeled with the second color (the “wild-type”). This could visualize the spread of the mutant in both uninfected and infected cells, and it could be tested whether these dynamics are more consistent with drift or with selection. HIV-1 could be a suitable experimental system [32]. This kind of experiment could then be repeated, but with a “wild-type”-infected cell population that has down-regulated the CD4 receptor, thus blocking entry of the “mutant” virus into wt-infected cells.

Following this initial spread, the dynamics of these mutants become more typical. Hence, the neutral mutant enters the phase of neutral drift, and the disadvantageous mutant experiences a selective disadvantage. In either case, multiple infection reduces the probability that the mutant spreads stochastically through the virus population, and makes virus extinction more likely. This leads to the counter-intuitive result that multiple infection can promote the presence of neutral or disadvantageous mutants in the short term, but reduces the chances to find those mutants in the longer term. As described above, this can complicate the interpretation of experimental results.

While it is important to understand the spread dynamics of the mutants, the physiologically most relevant scenario assumes that mutant viruses are generated repeatedly from the wild-type virus population by mutational processes, and that the newly created mutants attempt to spread. For advantageous mutants, the overall effect tends to be that a higher infection multiplicity results in a faster invasion of the mutant: While the probability of mutant fixation does not depend significantly on infection multiplicity, the rate of mutant generation is faster for higher multiplicity. For neutral or disadvantageous mutants, however, there is a tradeoff. While the rate of mutant generation is accelerated at higher infection multiplicity, the fixation probability of the generated mutant declines with higher multiplicity. The overall effect is a reduced rate of mutant invasion at high infection multiplicities, although for moderate multiplicities, the rate of mutant invasion can be faster than in the absence of multiple infection. These complex results indicate that multiplicity does not have a straightforward and consistent effect on the rate of mutant invasion. For example, the evolution of immune escape mutants in chronic infections that are controlled by ongoing immune responses is most likely accelerated by a higher infection multiplicity, since such mutants enjoy an instant fitness advantage. At the same time, however, other, equally important, evolutionary processes can be hampered at high multiplicities, such as the emergence of drug-resistant mutants before treatment initiation (standing genetic variation). Such mutants typically have a certain selective disadvantage compared to drug-sensitive viruses in the absence of treatment [33]. According to these results, it is therefore not possible to say that conditions in which viruses replicate at higher infection multiplicities either favor or hamper evolutionary processes.

The relatively complex dependence of basic evolutionary processes on infection multiplicity form an important foundation for further explorations of viral evolution. The consequences of recombination/ reassortment, complementation, and inhibition between wild-type and mutant viruses within the same cell have to be viewed as occurring on top of the basic dynamics described here in order to successfully understand the effect of these more involved interactions on evolutionary outcome.

